# *SvFUL2*, an A-class MADS-box transcription factor, is necessary for inflorescence determinacy in model panicoid cereal, *Setaria viridis*

**DOI:** 10.1101/2020.07.31.224378

**Authors:** Jiani Yang, Edoardo Bertolini, Max Braud, Jesus Preciado, Adriana Chepote, Hui Jiang, Andrea L. Eveland

**Affiliations:** Donald Danforth Plant Science Center, Saint Louis, MO, 63132, USA; National Science Foundation Research Experiences in Plant Science at the Danforth Center

## Abstract

Inflorescence architecture in cereal crops directly impacts yield potential through regulation of seed number and harvesting ability. Extensive architectural diversity found in inflorescences of grass species is due to spatial and temporal activity and determinacy of meristems, which control the number and arrangement of branches and flowers, and underlie plasticity. Timing of the floral transition is also intimately associated with inflorescence development and architecture, yet little is known about the intersecting pathways and how they are rewired during development. Here, we show that a single mutation in a gene encoding an AP1 A-class MADS-box transcription factor significantly delays flowering time and disrupts multiple levels of meristem determinacy in panicles of the C_4_ model panicoid grass, *Setaria viridis*. Previous reports of A-class genes in cereals have revealed extensive functional redundancy, and in panicoid grasses, no associated inflorescence phenotypes have been described. In *S. viridis*, perturbation of *SvFul2*, both through chemical mutagenesis and CRISPR/Cas9-based gene editing, converted a normally determinate inflorescence habit to an indeterminate one, and also repressed determinacy in axillary branch and floral meristems. Our analysis of gene networks connected to disruption of *SvFul2* identified regulatory hubs at the intersect of floral transition and inflorescence determinacy, providing insights into the optimization of cereal crop architecture.

## Introduction

Inflorescence structure determines fruit, seed, and pollen production, which are critical for reproductive success of plants and global food security. During the shift from vegetative to reproductive growth, the indeterminate shoot apical meristem (SAM), which patterns the vegetative organs, transitions to an inflorescence meristem (IM). Like the SAM, the IM continues indeterminate growth but instead, leaf growth is suppressed and axillary meristems (AMs) grow out into reproductive organs on its flanks. In eudicot systems such as *Arabidopsis thaliana*, the IM directly lays down floral meristems (FMs), which produce flowers. In grasses, FMs are borne from spikelet meristems (SMs) either directly from the IM as in wheat and barley, or after a series of AM branching events such as in maize and sorghum. Eventually AMs acquire SM identity and terminate in a spikelet, the central unit of the grass inflorescence, housing one to several flowers that bear grain. Variation in activity and determinacy of AMs and SMs in grasses allows for the wide diversity of inflorescence branching patterns (Tanaka et al., 2013; Whipple, 2017; Bommert and Whipple, 2018).

Inflorescence architecture is also shaped by the activity and determinacy of the IM. In certain cereals such as rice, barley and maize, the IM is indeterminate and continues meristematic activity, laying down lateral structures until it ceases growth. Alternatively, in wheat and sorghum, the IM takes on a determinate fate and produces a defined number of AMs before terminating in a spikelet. IM determinacy has been linked to flowering time through the action of multiple common regulators, which also affect branching patterns in the inflorescence (Danilevskaya et al., 2010; Li et al., 2019; Liu et al., 2019a). A weak flowering signal tends to delay meristem determinacy in the inflorescence, allowing for increased branch outgrowth and higher order branch initiation (McSteen et al., 2000; Endo-Higashi and Izawa, 2011; Boden et al., 2015).

Much of what we know about the molecular underpinnings of IM determinacy comes from Arabidopsis, which produces an indeterminate inflorescence. In Arabidopsis, indeterminacy in the IM is maintained by the antagonistic relationship between *TERMINAL FLOWER 1* (*TFL1*) and floral identity genes, *LEAFY* (*LFY*), *APETALA1* (*AP1*), and *CAULIFLOWER* (*CAL*) (Piñeiro and Coupland, 1998; Liljegren et al., 1999; Serrano-Mislata et al., 2017). *AP1* and *CAL* belong to the euAP1 subclade of the *AP1/FUL* (*FRUITFUL*)*-like MADS box* gene family and are key players in controlling flowering time and AM determinacy (Kempin et al., 1995; Alvarez-Buylla et al., 2006). *TFL1* expresses in the central region of the IM and prevents it from acquiring FM identity by suppressing floral identity genes (Weigel et al., 1992; Bradley et al., 1997; Benlloch et al., 2007). Loss of *TFL1* function results in the mis-expression of *AP1* and *LFY* in the IM, causing a terminal flower(s) to form in place of the indeterminate meristem, early flowering, and enhanced determinacy of lateral branches (Shannon and Meeks-Wagner, 1991; Alvarez et al., 1992). Alternatively, mutations in *AP1* and *LFY* genes result in production of indeterminate lateral shoots, which typically develop determinate FMs and have delayed flowering (Irish and Sussex, 1990; Schultz and Haughn, 1991; Huala and Sussex, 1992; Weigel et al., 1992; Bowman et al., 1993; Schultz and Haughn, 1993).

The regulatory modules that control inflorescence growth habit are somewhat conserved between eudicots and grasses. In maize and rice, *TFL1-like* genes delay flowering time and prolong the indeterminate status of the developing inflorescence (Nakagawa et al., 2002; Danilevskaya et al., 2010; Kaneko-Suzuki et al., 2018). In rice, *AP1/FUL-like* genes have overlapping roles in flowering time (Kobayashi et al., 2012). Over-expression of *OsMADS14, OsMADS15*, or *OsMADS18* all result in early flowering phenotypes (Jeon et al., 2000; Fornara et al., 2004; Lu et al., 2012), and in the case of *OsMADS15*, reduced panicle size and branch number (Lu et al., 2012). In winter wheat and barley varieties, expression of *VERNALIZATION 1* (*VRN1)*, an *AP1/FUL-like* gene, has been well-characterized as an early signal in promoting timely vegetative-to-reproductive transition in response to vernalization (Yan et al., 2003; Preston and Kellogg, 2008; Li et al., 2019). Expression of *FUL2* and *FUL3* genes in wheat are also induced by vernalization to promote flowering (Chen and Dubcovsky, 2012; Li et al., 2019). A recent study revealed that *AP1/FUL-like* genes in wheat and the genetic interactions among them contribute to maintenance of IM and SM determinacy, as well as flowering time (Li et al., 2019). Loss-of-function in both *VRN1* and *FUL2* genes converted the normally determinate IM of the wheat spike to an indeterminate habit, and also enhanced indeterminacy in primary AMs. Introduction of a single functional copy of either *VRN1* or *FUL2* reverted the *vrn-null; ful1-null* mutant IM back to a determinate habit (Li et al., 2019).

While evidence across the plant kingdom supports conserved roles for *AP1/FUL-like* genes in floral transition and inflorescence architecture, to date there have been no inflorescence phenotypes described for loss-of-function A-class genes in the subfamily *Panicoideae*, which includes agronomically important crops such as maize, sorghum, and sugarcane. This is likely due to functional redundancy (Litt and Irish, 2003; Preston and Kellogg, 2007). In this study, we show that a single loss-of-function mutation in an *AP1/FUL-like* gene in model panicoid grass, *Setaria viridis* (green foxtail), is sufficient to confer both strong flowering time and inflorescence determinacy phenotypes despite its overlapping expression pattern with three closely related paralogs. *S. viridis* is a weedy, C_4_ species that has demonstrated promise as a model system for elucidating molecular mechanisms in panicoid crops (Li and Brutnell, 2011; Huang et al., 2017; Yang et al., 2018). It also represents a key evolutionary node between domesticated and undomesticated grasses. Like wheat, *S. viridis* produces a determinate inflorescence that terminates in a spikelet, but AMs undergo multiple orders of branching (Doust and Kellogg, 2002a; Zhu et al., 2018). We isolated the *Svful2* mutant in a genetic screen, which displayed a ‘barrel’-like panicle morphology due to enhanced indeterminacy in AMs. The determinate IM was also converted to an indeterminate habit resembling a maize ear. Further investigation of *Svful2* loss-of-function at the molecular level using genomics approaches revealed regulatory modules that link floral transition and inflorescence determinacy pathways through interactions among MADS-box TFs and several other developmental regulators. This mutant and the analyses presented here, provide insights into the complex interface of flowering time and inflorescence development, and potential targets for fine-tuning inflorescence ideotypes in cereal crops.

## Results

### Characterization of the *barrel 1* (*brl1*) mutant in *Setaria viridis*

In a forward genetics screen of ∼3000 N-methylurea (NMU) mutagenized M2 families of *S. viridis (Huang et al*., *2017; Yang et al*., *2018)*, we isolated the *barrel1* (*brl1*) mutant, named for its abnormal, barrel-shaped panicle. Compared with mature panicles of the wild-type mutagenized reference line (A10.1), mutant panicles were shorter and thicker and appeared more brachy (Fig. 1, A and B; Table 1). Mutant plants were shorter in stature and produced tillers with more leaves. In addition to morphological defects, flowering time was obviously delayed in *brl1* mutants. To test the effect of different photoperiods on floral transition, we examined flowering time of the mutant compared to control plants under both short-day (SD, 12 hours light /12 hours dark) and long-day (LD, 16 hours light / 8 hours dark) conditions. Under SD conditions, which typically promote flowering in *S. viridis* (Doust et al., 2017), *brl1* mutant panicles emerged approximately six days later (avg. 29.20 Days After Sowing (DAS)) than those of wild-type (avg. 22.94 DAS; Fig. 1C). Under LD conditions, flowering time in *brl1* mutant (avg. 31.21 DAS) and A10.1 wild-type plants (average at 28.44 DAS) was delayed compared to under SDs, but *brl1* mutants still flowered significantly later than wild type (avg three days, Fig. 1D; Table. S1).

**Figure 1.**
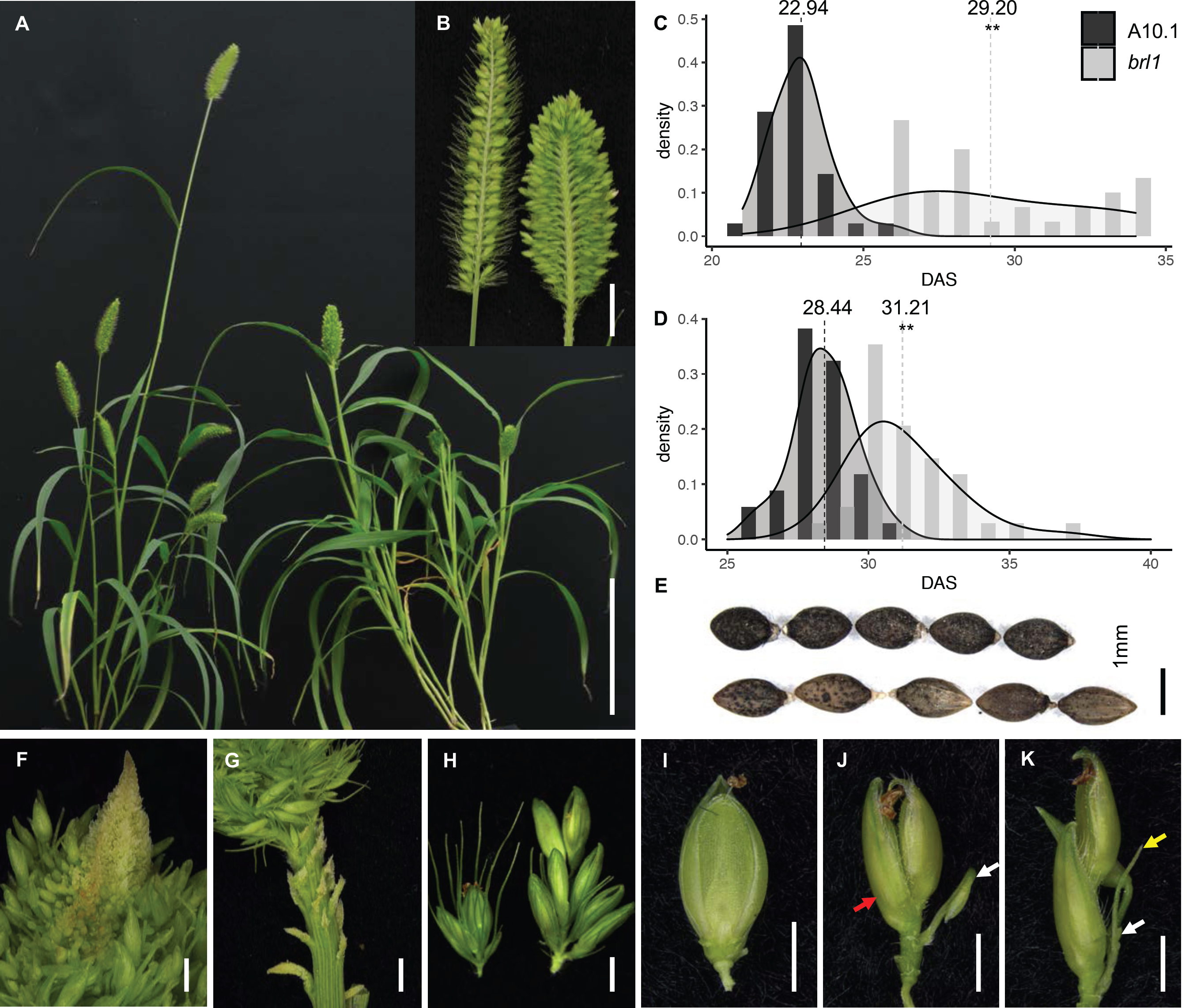
Characterization of the *brl1* mutant phenotypes. **(A)** Plant morphology of the wild-type A10.1(left) and the *brl1* mutant (right). Scale bar = 10 cm. **(B)** Compared to the wild-type (left), *brl1* mutant (right) panicles are shorter and wider and primary branches are packed more densely. Primary branches were removed from one side of the panicle for a longitudinal view. Scale bar = 1 cm. Under SD **(C**) and LD **(D**) conditions, panicles of the *brl1* mutant emerged significantly later than those of wildtype. **: *p*-value < 0.01. **(E)** Compared to wild-type (upper panel), the *brl1* mutant (lower panel) produced longer and narrower seeds. **(F)** In *brl1* mutant panicles, the IM appeared indeterminant with continual production of primary branches. Scale bar = 1 mm. **(G)** Rudimentary primary branches were visible at the base of mature panicles of the *brl1* mutant. Scale bar = 1 mm. **(H)** Primary branches in the *brl1* mutant panicles (right) are markedly longer than those of A10.1 (left). Scale bar = 1 mm. Examples of phenotypes in *brl1* mutant spikelets that lost SM maintenance (**J and K**), including aberrant development of the lower floret (red arrow, **J**) or production of additional bristles (yellow arrow, **K**) and spikelets (white arrow, **J and K**) within a spikelet, compared to A10.1 (**I**). Glumes were removed in **I, J and K** for better view. Scale bars = 1 mm.

Previous studies in *S. viridis* showed that flowering time impacted both plant architecture and biomass (Doust, 2017). Under both LD and SD conditions, plant height and panicle length of *brl1* mutants were significantly shorter than wild-type plants at maturity. Under SD conditions, above-ground dry weight was increased in mutants compared to wild-type, largely due to biomass of vegetative tissue (leaves and stems; Fig. S1). In LDs, above-ground dry weight of *brl1* mutants was comparable to wild-type, however we still observed a significant increase in dry weight of vegetative tissue (leaves and stems) in the mutant (Fig. S1). Seed shape and size were also different with the mutant seeds being longer and narrower than those of wild-type (Fig. 1E; Table 1).

Examination of the inflorescence morphology revealed that *brl1* mutants displayed various levels of indeterminacy. At the tip of the panicle, the IM appeared indeterminate in mutants, and newly formed branch meristems (BMs) were still visible at maturity (Fig. 1F). At the base of the mutant panicle, rudimentary primary branches were observed, which were not found in wild type (Fig. 1G). Primary branches were longer in *brl1* mutants and the panicle rachis was clearly thicker (Fig. 1H; Supplemental Fig. S1A). Bristles, which are modified branches paired with spikelets in *Setaria sp*., did not elongate to the length of wild-type bristles, and so were largely found buried under spikelets (Fig. 1H). Development of spikelets and flowers was also affected in *brl1* mutants, but phenotypes showed low penetrance with varied severities of indeterminacy. For example, approximately 17% of *brl1* mutant panicles produced additional flowers, bristles and/or spikelets within spikelets compared to the typical one flower per spikelet in wild-type (Fig. 1, I-K). The lemma and palea of mutant flowers were more elongated in the mutant and were more rigid, which is likely contributing to the elongated seed shape (Supplemental Fig. S1 B and C).

### *brl1* mutants show loss of determinacy in various stages of inflorescence development

We used Scanning Electron Microscopy (SEM, Fig. 2) to compare the developmental progression of inflorescence primordia from the *brl1* mutant with that of wild-type *S. viridis*. By 11 DAS, the vegetative SAM of wild-type plants had finished transitioning to the reproductive IM, as the first primary BMs were initiated on its flanks (Fig. 2A). In the *brl1* mutant, the vegetative-to-reproductive transition was delayed to 15 DAS (Fig. 2B), consistent with its late flowering phenotype (Fig. 1C). After the transition, wild-type inflorescences initiated primary branches in a spiral pattern (Fig. 2C), and then secondary and tertiary axillary branches sequentially in a distichous pattern, as previously described (Fig. 2E) ((Doust and Kellogg, 2002a; Yang et al., 2018; Zhu et al., 2018)). The *brl1* IM was elongated compared to that of wild-type (Fig. 2D, Supplemental Fig. S1 D and E), and this appeared to enable capacity for increased initiation of primary and higher order branches (Fig. 2, D and F), consistent with the mature panicle phenotype. By 17 DAS the wild-type IM had become determinate and terminated as a spikelet (Fig. 2G). BMs then began to differentiate from the tip of the inflorescence primordium into either a spikelet meristem (SM) or a sterile bristle, and this continued basipetally (Fig. 2G). Conversely, the IM of the *brl1* mutant remained indeterminate and continued to produce primary BMs at 21 DAS, where SMs and bristles began to differentiate towards the top of the inflorescence primordium (Fig. 2H). By the end of the developmental series analyzed by SEM, the *brl1* IM remained indeterminate, which is consistent with its mature phenotype in Fig. 1F.

**Figure 2.**
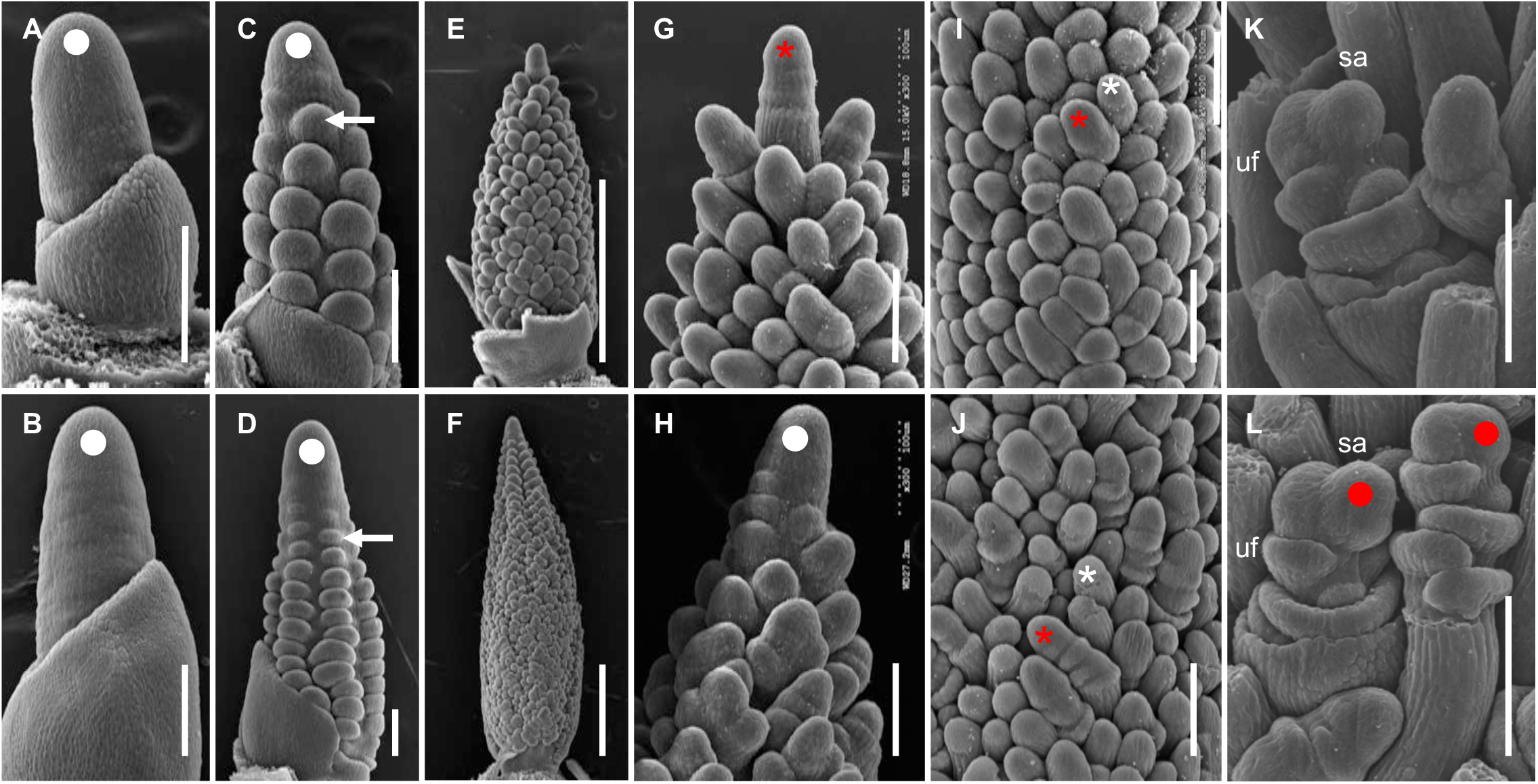
Morphological analysis of early inflorescence development in the *brl1* mutant by SEM. The transition from SAM to IM **(white dot)** in the *brl1* mutant was delayed to 15 DAS (**B**) compared to 11 DAS (**white dot**) in A10.1 (**A**). Branching capacities were increased in *brl1* panicles (**D and F**, 18 DAS and 20 DAS, respectively) compared to those of A10.1 (**C and E**, 12 DAS and 14 DAS, respectively). In A10.1, the 17 DAS IM ceased to produce new BMs and terminated as the first SM (**G, red asterisk**) and then BMs started to differentiate into SMs (**I, red asterisk**) and bristles (**I, white asterisk**) basipetally. However, at 21 DAS, the *brl1* mutant IM continued initiating primary branches at the inflorescence tip (**H, white dot**), even after BMs acquired SM (**J, red asterisk**) or bristle identities (**J, while asterisk**). Some *brl1* spikelets had abnormal outgrowth of meristems (**red dots**) in upper florets (**L**) compared to A10.1 spikelet **(K**). Scale bars = 100 µm in **A-D** and **G-L**. Scale bars = 500 µm in **E and F**. uf: upper floret and sa: spikelet axis.

While differentiation of SMs and bristles appeared normal in the mutant (Fig. 2, I and J), the onset was delayed compared to wild-type and after additional rounds of higher order branching (Fig. 2F). SMs developed similarly in *brl1* mutants and wild-type, initiating glumes and upper and lower FMs; the upper floret typically develops into a perfect flower with lemma, palea, anther, and carpel and the lower floret aborts (Doust and Kellogg, 2002b; Yang et al., 2018). In some cases, we observed aberrant meristematic outgrowths in *brl1* FMs (Fig. 2L) which may explain our observations of additional spikelets and bristles within some spikelets (Fig. 1, I and J). Our SEM analysis showed that a determinacy program was delayed in the IM, BMs, and SMs of *brl1* mutants.

### *The brl1 locus* encodes *SvFUL2*, a MIKC-type MADS-box transcription factor

F2 populations were generated from a cross between the *brl1* mutant and the parental line, A10.1. Wild-type and barrel-like panicle phenotypes segregated with the expected Mendelian 3:1 ratio (139:48; P (χ2, 1 d.f.) = 0.83), which indicated that *brl1* is a single locus recessive allele. To map the *brl1* locus, Bulk Segregant Analysis (BSA) was performed (Michelmore et al. 1991; Schneeberger 2014) with a pool of DNA from 30 *brl1* mutant individuals from the segregating F2 population that was sequenced to ∼17 x coverage (65.6M reads). Reads were aligned to the A10.1 reference genome (phytozome.jgi.doe.gov; v1.1; (Mamidi et al., 2020)) and single nucleotide polymorphisms (SNPs) were called by GATK (3.5-0-g36282e4). Unlike our previous experience with mapping by BSA in this population (Yang et al., 2018), we did not resolve a clear peak in a genomic region of low variability, likely due to lower read coverage. However, several high-confidence, nonsynonymous SNPs were identified and supported by high observed allele frequency (Supplemental Fig S2, Supplemental Data Set 1). One candidate SNP disrupted the start codon of Sevir.2G006400, a MIKC-type MADS-box gene, and was supported by whole genome sequencing of the *brl1* mutant. Sevir.2G006400 had previously been annotated as *SvFul2* based on phylogenetics and evolutionary developmental analyses (Preston et al., 2009; Zhu et al., 2018).

We designed a dCAPs marker specific for this SNP and genotyped over 200 segregating F2 individuals. Our genotyping results showed that this SNP co-segregated with the barrel panicle phenotype at 100% (Supplemental Fig. S3). We also used RT-PCR to test whether expression of Sevir.2G006400 was disrupted in *brl1* mutant inflorescence primordia. Results showed that at an early stage of inflorescence development, Sevir.2G006400 was expressed at a lower level in *brl1* mutants compared with A10.1 (Supplemental Fig. S4).

*SvFul2* encodes the ortholog of *OsMADS15* in rice, an *AP1/FUL-like* MADS-box gene in the MIKC-type subfamily. Consistent with previous phylogenetic studies (Wu et al., 2017; Li et al., 2019), our phylogenetic analysis of *AP1/FUL-lik*e MADS-box genes from *S. viridis* as well as Arabidopsis, rice, wheat, maize, and sorghum showed that *SvFul2* was located in the *FUL2* subclade along with three copies of wheat *Ful2s*, rice *OsMADS15*, and maize *zap1* and *zmm3* (Fig. 3A). *SvFul2* is more closely related to *SvFul1* (Sevir.9G087300) in the *FUL1* subclade, which includes wheat *VRN1s*, rice *OsMADS14*, maize *zmm15* and *zmmads4. SvFul3* (Sevir.2G393300) and *SvFul4* (Sevir.3G374401) are located in the *FUL3* and *FUL4* subclades, respectively. By examining a previously generated transcriptomics resource across six sequential stages of early *S. viridis* inflorescence development (Zhu et al., 2018), we found that *SvFul1, SvFul2*, and *SvFul3* shared similar spatiotemporal expression patterns, increasing during branching and then decreasing during floral development with a small drop during spikelet specification (Fig. 3B). *SvFul1* was expressed highest at 10 and 12 DAS and *SvFul2* expressed more at later stages, which indicate the two may have different functions. Comparatively, *SvFul4* was expressed lower throughout inflorescence development, its expression gradually decreasing after the reproductive transition.

**Figure 3.**
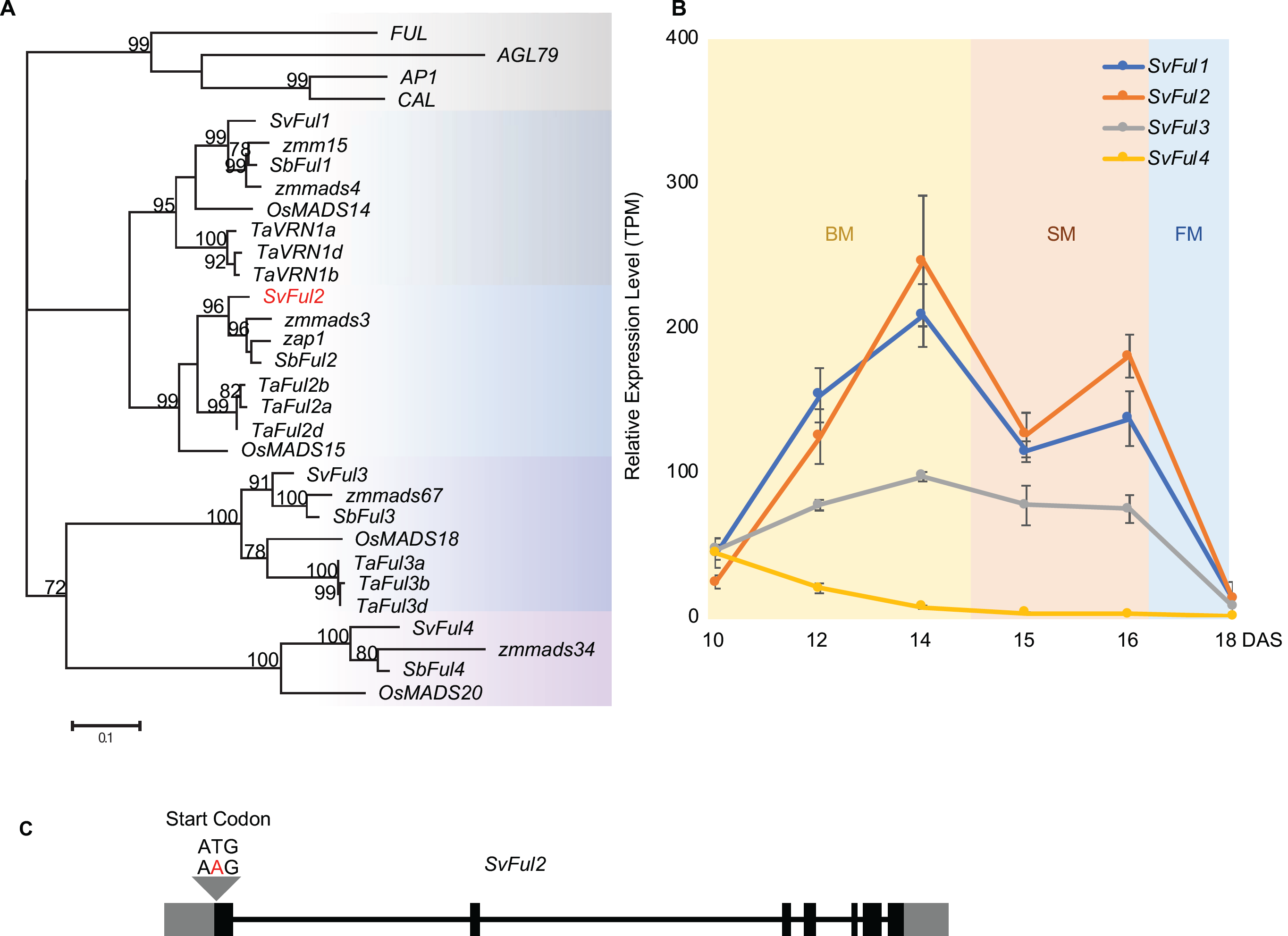
Phylogeny of *AP1/FUL-like* MADS-box genes in grasses and Arabidopsis and their expression profiles in *S. viridis* inflorescence development. **(A)** Phylogenetic analysis of *AP1/FUL-like* MADS-box genes was based on protein coding sequence. *SvFul2* is highlighted in red. **(B)** Expression patterns of four *S. viridis AP1/FUL-like* MADS-box genes across six stages of early inflorescence development based on the resource described in (Zhu et al., 2018). Error bars indicate standard errors of three to four biological replicates. BM, branch meristem; SM, spikelet meristem; and FM, floral meristem. **(C)** Exon-intron structure of the *SvFul2* gene consists of seven exons (**solid rectangles**) and six introns (**horizontal line**). The 5′ and 3′ untranslated regions are shown as **grey rectangles. Grey triangle** indicates the location of the SNP that disrupts the start codon within the *SvFul2* gene.

### Gene editing of *SvFul2* validates the mutant phenotype in *S. viridis*

To validate that Sevir.2G006400 (*SvFul2*) is responsible for the observed phenotypes of the *brl1* mutant, we used genome editing. A CRISPR/Cas9 construct was designed containing two guide (g)RNAs that specifically targeted the first exon of *SvFul2* in the highly transformable *S. viridis* accession, ME034 (Fig. 4A; (Acharya et al., 2017; Van Eck, 2018). In the T1 generation, individual plants carrying a homozygous 540 bp deletion in the 1st exon of *SvFul2* were selected (Fig. 4A, Supplemental Fig S2). We called this genotype *SvFul2_KO*. These were moved forward to generation T2 where they were then outcrossed to ME034 and then selfed to select Cas9-free *SvFul2_KO* plants for phenotyping. *SvFul2_KO* plants displayed phenotypes consistent with those of the *brl1* mutant (Fig. 4B-G). Compared with ME034 normal plants, *SvFul2_KOs* were shorter and branchy with more leaves, and panicles displayed increased densities of longer primary branches (Fig. 4C and F). As observed in *brl1* mutants, panicles of *SvFul2_KO*s took on an indeterminate growth habit (Fig. 4C-F). Flowering time was also delayed in the *SvFul2_KOs* (avg. 20.76 DAS) compared to the ME034 wild-type siblings (avg. 17.87 DAS) by approximately three days (Fig. 4G). The ME034 accession flowers earlier than A10.1, consistent with the shift in flowering time shown here.

**Figure 4.**
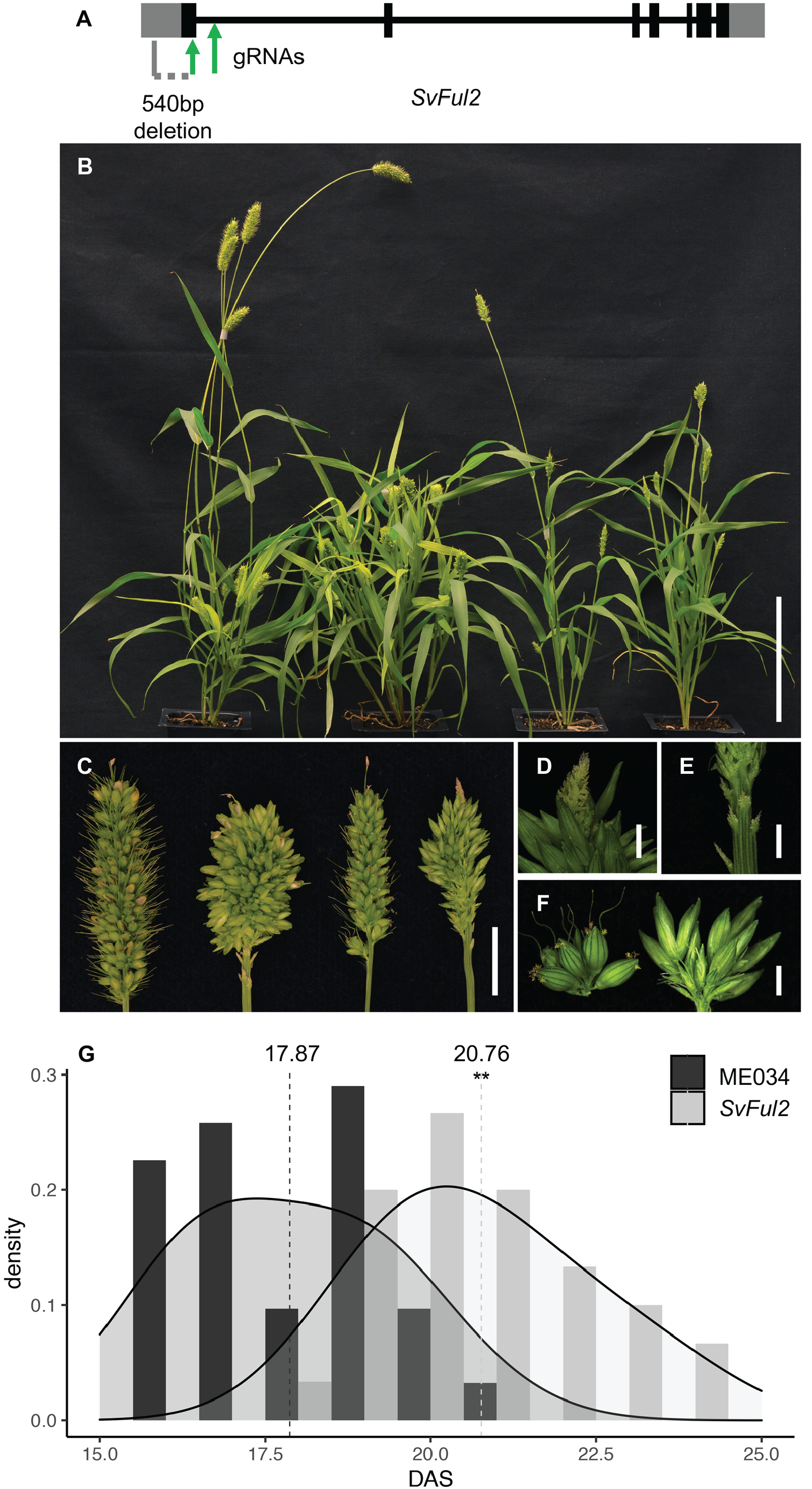
CRISPR/Cas9-based gene editing of *SvFul2* phenocopied the *brl1* mutant. **(A)** Schematic diagram of the *SvFul2* gene model showing locations of the two (g)RNAs target sites (**green arrows**) and the 540bp deletion region (**grey dash line**) in *Svful2_KO* plants. Plant morphology (**B**) and inflorescence structure (**C**) of the *Svful2_KO* mutant phenocopied that of the *brl1* mutant. From left to right: A10.1, *brl1*, ME034, and *Svful2_KO* individuals. Scale bars equal 10cm and 1cm in A and B, respectively. Indeterminate IM (**D**), underdeveloped primary branches at the panicle base (**E**), and longer primary branches (**F**) were also observed in *Svful2_KO* panicles. Scale bars equal 1mm. (**G**) Under SD conditions, panicle emergence day of *Svful2_KO* is significantly delayed compared to ME034. **: *p*-value < 0.01.

We further tested the allelic relationship between the *brl1* mutant in the A10.1 background and *SvFul2_KO* in ME034 by crossing *brl1* with *SvFul2_KO*. The resulting bi-allelic F1 individuals displayed a barrel phenotype and failed to complement, indicating that they are allelic (Fig. 5). Taken together, our analyses support *SvFul2*, Sevir.2G006400, as the locus responsible for the *brl1* mutant phenotypes in *S. viridis*.

**Figure 5.**
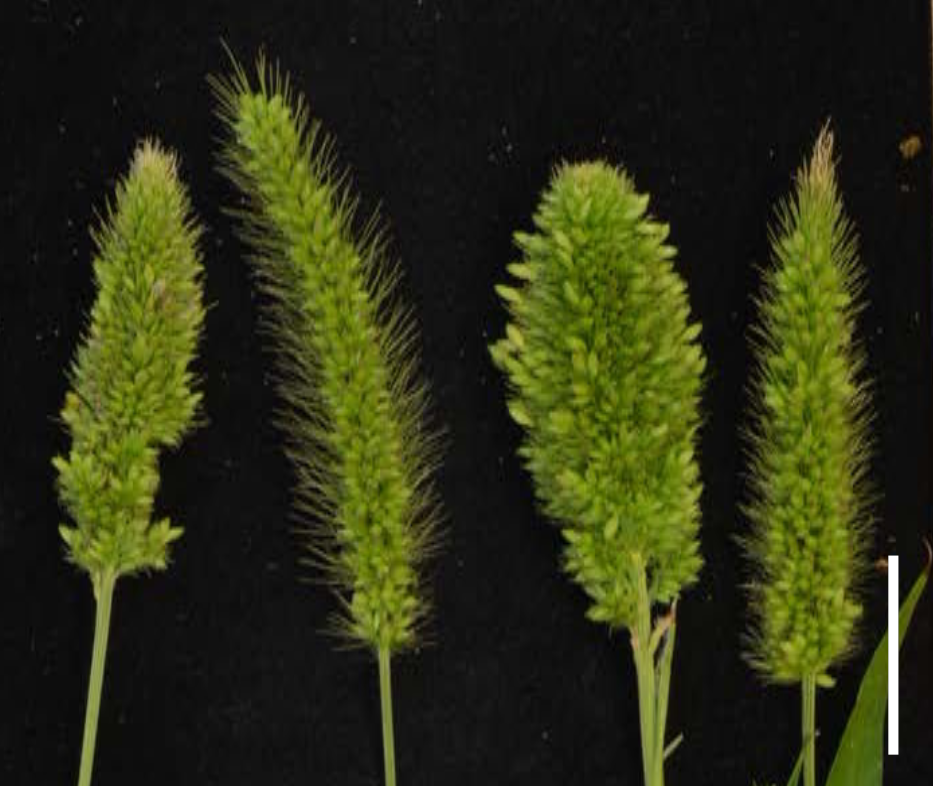
Allelism test between *brl1* and *Svful2_KO* fails to complement the mutant phenotype. The *brl1* mutant was crossed to *Svful2_KO* and the resulting bi-allelic F1 was compared with the F1 plants from crosses between A10.1 and *Svful2_KO, brl1* mutant and A10.1. From left to right: panicles from *brl1*/*Svful2_KO* F1, A10.1/*Svful2_KO* F1, *brl1* mutant, and A10.1. Scale bar equals 2cm.

### Loss of *SvFul2* function alters expression of flowering and meristem determinacy pathways

To determine the molecular mechanisms underlying the complex phenotypes of the *Svful2* mutant, we used RNA-seq to profile gene expression in mutant inflorescence primordia across three key developmental transitions and compared them to equivalent stages in wild-type controls: right before (stage 1) and after (stage 2) the floral transition, and during the initiation of spikelet specification (stage 3). Here, we expect to capture transcriptional changes related to both differences in flowering time and meristem determinacy. For each stage, we profiled four biological replicates, each consisting of pooled, hand-dissected inflorescence primordia. Differential expression was determined using DESeq2 (1.22.2). Our analysis found 382, 2,584, and 2,035 differentially expressed genes (DEGs) at stages 1, 2, and 3, respectively. Based on Principal Components Analysis (PCA), we observed fewer differences in the mutant transcriptome at stage 1, suggesting that the main influence of *SvFul2* on inflorescence development begins once the SAM has initiated transition to the IM (Fig. 6A). We also observed dynamic shifts in DEGs among the three stages; only 33 DE genes were shared across all three stages, and 149, 451, and 68 were shared between stage 1 and 2, stage 2 and 3, and stage 1 and 3, respectively (Fig. 6B). This suggests that *SvFul2* potentially modulates different target genes in various spatiotemporal contexts.

**Figure 6.**
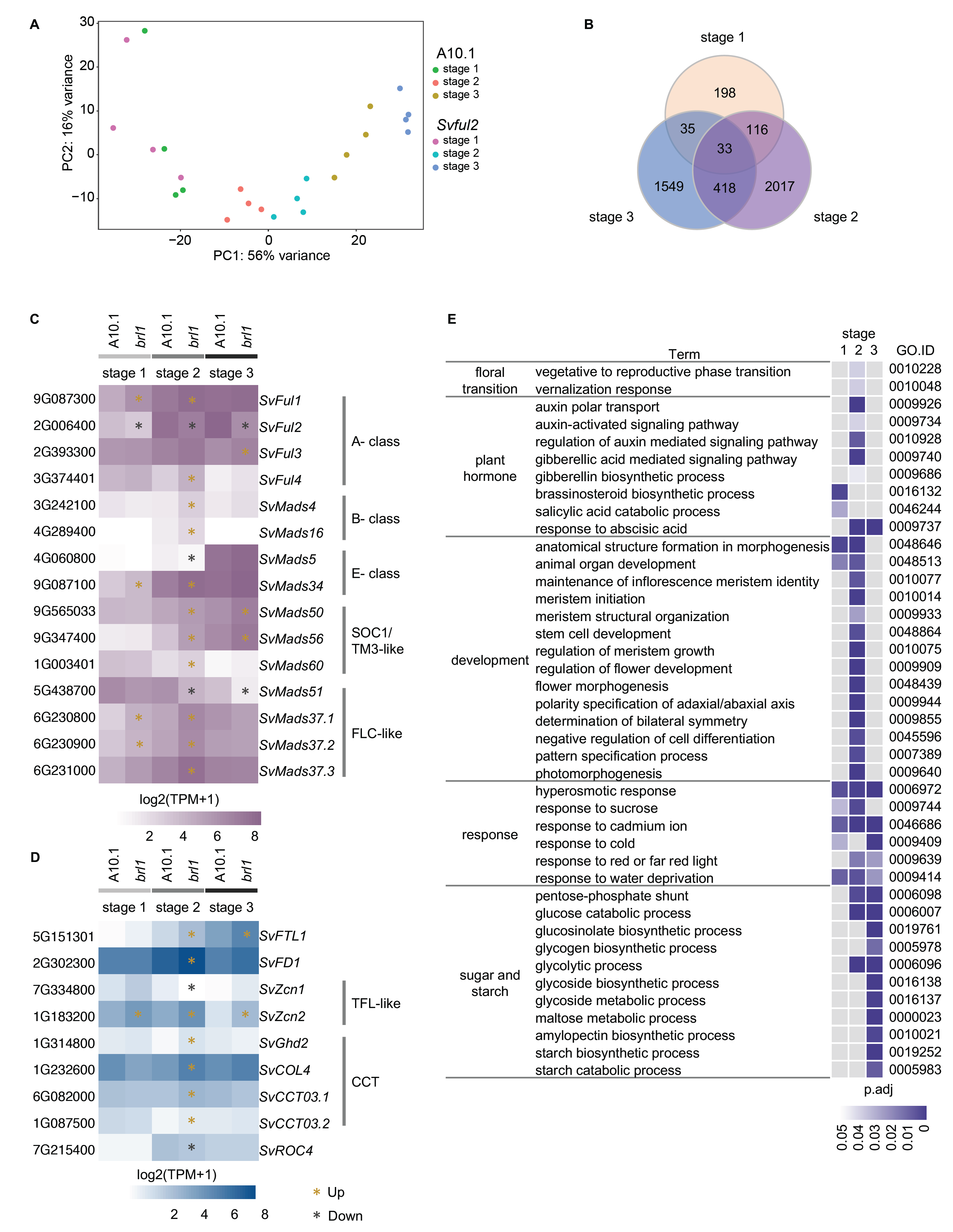
Transcriptional changes of *Svful2* mutants across three stages of inflorescence development. **(A)** PCA analysis showed that biological replicates were well-correlated with each other and that PC1 (explaining 56% of the variance) was associated with developmental stage. Loss-of-function in *Svful2* resulted in fewer transcriptional changes prior to the floral transition with larger changes between genotypes becoming evident after. **(B)** Differential expressed genes showed dynamic transcriptional changes in *Svful2* mutants at three stages of inflorescence development. Among differentially expressed genes were several encoding MADS-box TFs (**C**) and known regulators of flowering time (**D**). TPM values were Log2 transformed to generate heatmaps. Yellow and black asterisks indicate up- and down-regulated DEGs (FDR < 0.05), respectively. (**E**) Subsets of GO terms that were overrepresented among DEGs at each of the three developmental stages. p.adj < 0.05.

As expected, the *SvFul2* gene itself was significantly down-regulated in mutant inflorescences at all three stages (Fig. 6C). The other three *S. viridis AP1/FUL-like* genes were significantly up-regulated in the mutant, suggesting that the four *AP1/FUL-like* genes may provide some level of functional compensation during inflorescence development. Two B-class genes, *SvMads16* (*AP3*) and *SvMads4* (*PISTILLATA*), which in grasses are typically expressed at low levels prior to floral organ development (Whipple et al., 2004), were up-regulated in *Svful2* stage 2 inflorescences (Fig. 6C). In addition, two E-class genes were differentially expressed in mutant inflorescences: *SvMads34* was up-regulated at stages 1 and 2, while *SvMads5* was down-regulated at stage 2 (Fig. 6C). In rice, *OsMADS34* coordinates with *AP1/FUL-like* genes, and physically interacts with some of them, to specify inflorescence meristem identity (Kobayashi et al., 2012). In general, E-class genes play partially redundant roles in specifying floral organ identities via protein-protein interactions with other MADS box proteins (Pelaz et al., 2000; Honma and Goto, 2001; Theißen and Saedler, 2001; Ditta et al., 2004).

Since the transition to reproductive growth is delayed in *Svful2* mutants, we expected to see changes in genes and pathways associated with flowering time (Fig. 6D). Functional categories related to flowering time were overrepresented among DEGs post-transition at stage 2, including “vegetative to reproductive phase transition of meristem” (GO:0010228; p.adj = 3.75e^-02^) and “vernalization response” (GO:0010048; p.adj = 3.68e^-02^) (Fig. 6E). Homologs of well characterized genes known to regulate flowering in other species were differentially expressed (Fig. 6D). For example, the *S. viridis* ortholog of *OsFTL1* (Sevir.5G151301, *SvFtl1*) encoding florigen (FT protein) and the ortholog of rice *OsFD1* (Sevir.2G302300, *SvFd1*), were up-regulated at stage 2. In Arabidopsis, *FD* is repressed by AP1 (Kaufmann et al., 2010). In addition, members of *FLC-like* and *TM3/SOC1-like* MIKC-type MADS-box genes and *CO (CONSTANS)-like* genes, which are also implicated in floral transition (Zhang et al., 2015; Huang et al., 2018), were among DEGs up-regulated in *Svful2* mutants at stage 2. Certain *S. viridis* orthologs of *zea mays centroradialis* (*zcn*) genes encoding FT homologs were differentially expressed, including *SvZcn2* (Sevir.1G183200), which is phylogenetically closest to Arabidopsis *TFL1* (Danilevskaya et al., 2008; Danilevskaya et al., 2010), was up-regulated in *Svful2* mutants at all three stages. In Arabidopsis, AP1 and TFL1 act antagonistically to repress each other’s expression to modulate flowering time and IM determinacy (Shannon and Meeks-Wagner, 1991; Alvarez et al., 1992; Kaufmann et al., 2010).

Consistent with defects in meristem determinacy, DEGs in *Svful2* mutants were enriched for functions related to meristem development with overrepresented GO terms such as “meristem initiation” (GO:0010014; p.adj = 7.63e^-04^) and “stem cell development” (GO:0048864; p.adj = 0.0046) (Fig. 6E). Up-regulated DEGs included homologs of *AP2-like* genes in maize known to suppress indeterminate growth in the SM; *SvIds1* (*indeterminate spikelet1*, Sevir.9G034800) and *SvSid1* (*sister of indeterminate spikelet1*, Sevir.2G093800; Supplemental Fig S6) (Chuck et al., 1998; Chuck et al., 2007; Chuck et al., 2008). The homolog of rice *OsMTF1* (*MOTHER OF FT AND TFL1*), which represses SM identity, was also up-regulated in *Svful2*, and the ortholog of maize *ramosa2* (*SvRa2;* Sevir.5G116100), which functions to promote meristem determinacy (Bortiri et al., 2006), was down-regulated.

As a consequence of increased meristem indeterminacy, *Svful2* mutant inflorescences branch more. We also found that genes associated with “anatomical structure formation involved in morphogenesis” (GO:0048646) were overrepresented among DEGs at stages 1 (p.adj = 0.0027) and 2 (p.adj = 3.56e^-08^; Fig. 6E), consistent with enhanced expression of genes involved in organogenesis. Among this functional class were orthologs of known genes that specify abaxial cell fate, e.g., Sevir.1G255800 (*SvYab15*) and *SvMwp1* (*milkweed pod1*, Sevir.6G158800) (Candela et al., 2008), and adaxial cell fate, e.g., *SvRoc5* (*Rice outermost cell-specific gene5*, Sevir.5G077800) and *SvPhb* (*PHABULOSA*, Sevir.9G157300) (McConnell et al., 2001; Juarez et al., 2004; Zou et al., 2011), which were up-regulated in the mutant (Supplemental Fig S6). Alternatively, the ortholog of rice *DWARF3, SvD3* (Sevir.4G068300), which functions in suppression of branching through the strigolactone signaling pathway (Zhou et al., 2013), was down-regulated. The major transcriptome changes observed in loss-of-function mutants at stage 2 reflect a core function for SvFUL2 in modulating the reproductive transition at the molecular level, but also how it links delayed flowering to suppression of meristem determinacy programs.

### Transcriptional rewiring by perturbation of *SvFul2* reveals sub-networks connecting reproductive transition and determinacy pathways

To further investigate how *SvFul2* connects within a larger gene network to regulate flowering time and meristem determinacy pathways, we used a computational strategy based on weighted gene co-expression network analysis and a random forest classifier to construct a gene regulatory network (GRN) representing normal inflorescence development in *S. viridis* (A10.1). Here, we integrated RNA-seq data from a previous study that captured precise stages of A10.1 inflorescence primordia spanning the IM transition to the development of floral organs (Zhu et al., 2018) with the staged wild-type data collected in this study. Using the WGCNA algorithm (Langfelder and Horvath, 2008) we clustered 26,758 genes into 27 co-expression modules (Fig. 7A; Supplemental Fig S7). Module eigengenes (expression pattern that best fits an individual module) were evaluated for their significant associations with four key developmental events represented in the network: the vegetative-to-reproductive transition (8 and 10 DAS), branching (11, 12, and 14 DAS), meristem determinacy (15-17 DAS), and flower development (18 DAS) (Fig. 7A). Within each module, we tested for enrichment of genes that were differentially expressed in the *Svful2* mutant, and found several that showed enrichment at certain stages of development (Fig. 7A). Among these, the module eigengene MEmagenta showed a strong positive correlation with the floral transition and a negative correlation with meristem determinacy (Fig. 7A-B). MEmagenta showed enrichment for DEGs in stages 1 and 2 (Fig. 7A). Alternatively, MEbrown showed a positive correlation with meristem determinacy, but was negatively correlated with the floral transition (Fig. 7A-B). *SvFul2* and *SvFul3* were both co-expressed in the brown module, along with 428 DEGs largely at stages 2 and 3.

**Figure 7.**
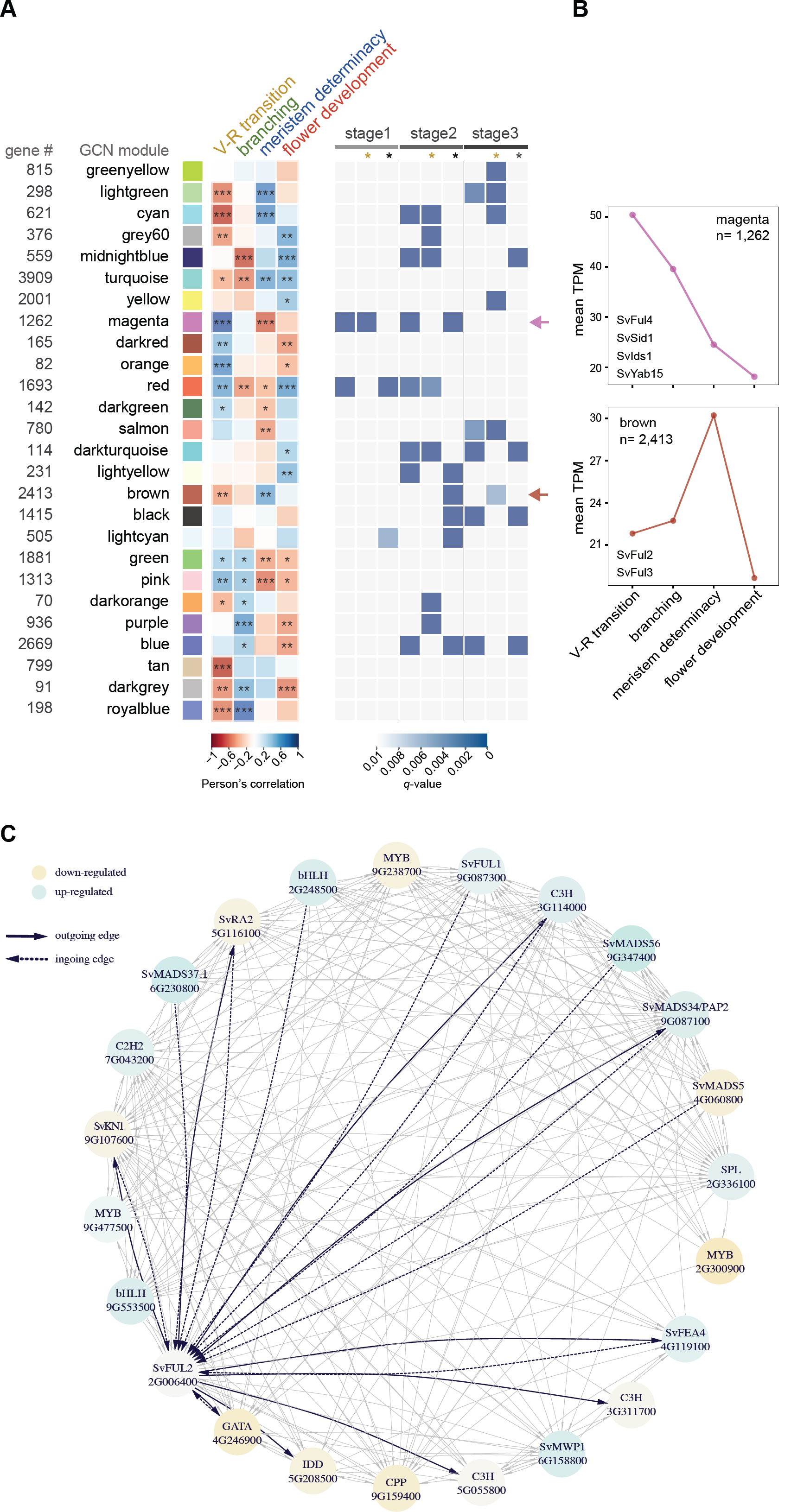
Network analysis reveals transcriptional rewiring in the *Svful2* mutants that link reproductive transition and determinacy pathways. **(A)** *(left)* Heatmap represents the WGCNA eigengene module association with key events during early inflorescence development in *S. viridis*. Network modules are represented and named with different colors based on the WCGNA default module annotation. Number of genes co-expressed in each module are indicated to the left. Student asymptotic *p*-value for the eigengene module association are indicated: ***, p.value<0.001; **, p.value<0.01, and *, p.value<0.05. *(right)* Heatmap represents enrichment of *Svful2* DEGs at each of the three developmental stages profiled in the mutant among the eigengene modules. Gold and black stars indicate up- and down-regulation, respectively, in the *Svful2* mutant compared to wild-type. **(B)** Expression trajectories of MEmagenta and MEbrown across *S. viridis* inflorescence development. Y-axis represents the average TPM of co-expressed genes in each module. **(C)** Subnetwork of predicted direct targets of SvFUL2 and its direct upstream regulators based on our GRN. Genes are represented as circles with edges linking specific regulators to their targets. Differential fold change of expression in the *Svful2* mutant background is represented by the colored scale. Darker colors represent higher |fold change|.

We also integrated the co-expression network with information derived from regulatory interactions among TFs and their putative targets based on the GENIE3 algorithm (Huynh-Thu et al., 2010). This complementary approach helped us to resolve the directionality and connectivity of important hub genes within the GRN. We selected TFs expressed in our dataset (n = 1,295) based on PlantTFDB (Jin et al., 2017) and used their trajectories in the network to derive points of connection with their potential target genes. Regulatory genes and their predicted targets were restricted based on information from differential expression analysis between wild-type and the *Svful2* mutant. We used the resulting regulatory framework to explore functional relationships between SvFUL2, its predicted direct targets, and predicted upstream regulators, particularly in the context of connecting floral transition and meristem determinacy. Our predictions indicated that SvFUL2 controls several co-expressed TFs previously implicated in developmental processes, and localized to modules that positively associated with branching/meristem determinacy and negatively associated with floral transition (Fig. 7D). Among these were SvRA2, and an INDETERMINATE DOMAIN (IDD) TF, several members of which have been involved in both the floral transition and determinacy, including the founding member from maize, *indeterminate 1* (*id1*) (Colasanti et al., 1998; Kozaki et al., 2004)).

Our analyses point to a possible feedback loop mechanism between SvFUL2 and SvRA2, where *SvFul2* is also a predicted direct target of SvRA2. We also observed putative feedback regulation between SvFUL2 and TFs encoded by the orthologs of maize *knotted 1* (*kn1;* TALE TF, Sevir.9G107600) and *fasciated ear 4* (*fea4;* bZIP TF, Sevir.4G119100), which promote meristem maintenance and differentiation, respectively (Bolduc et al., 2012; Pautler et al., 2015). Extensive feedback interactions among these developmental TFs could represent endogenous mechanisms for fine-tuning developmental processes during the floral transition and the development of the inflorescence. In maize, KN1 was shown by ChIP-seq to bind a maize paralog of *SvFul2, zap1* (Bolduc et al., 2012). Interestingly, several other MADS-box TFs were shown to directly target *SvFul2* based on predictions in our GRN: SvMADS37.1 (Sevir.6G230800), SvMADS56 (Sevir.9G347400), SvMADS5 (Sevir.4G060800), SvMADS34/PAP2 (Sevir.9G087100) and SvFUL1 (Fig. 7D). *SvMads34/Pap2* was also predicted to be a direct target of SvFUL2. These predictions are consistent with previous studies showing regulatory interactions among MADS-box TFs during the floral transition and inflorescence development. In rice, OsMADS37 and OsMADS56 have been functionally characterized as flowering time regulators. *OsMADS34*/*PAP2* has been shown to function redundantly with rice *AP1/FUL-like* genes to promote flowering and SM determinacy (Kobayashi et al., 2010; Kobayashi et al., 2012), and *OsMADS5* is involved in spikelet identity (Wu et al., 2018). Our analysis indicates that SvFUL2 may function as a core integrator at the interface of these closely linked developmental programs through feedback coordination with several other developmental TFs.

## Discussion

According to the classic ABCDE model of floral development in Arabidopsis, A-class *AP1/FUL-like* MADS-box genes have essential functions in modulating the floral transition and floral organ development (Irish and Sussex, 1990; Ferrándiz et al., 2000; Kaufmann et al., 2010). The roles of A-class genes have been the focus of extensive evolution and development studies (Litt and Irish, 2003; Preston and Kellogg, 2007), however relatively little is known about their functions in grasses. Given the complex branching patterns that arise post-floral transition and prior to flower development in grasses, it is expected that there would be some variation in function. In general, grasses show subtle variations on the traditional ABCDE model, however the underlying mechanisms are generally conserved (Ambrose et al., 2000; Whipple et al., 2007). Functional redundancy among A-class genes is widespread in grasses, and only recently have simultaneous perturbations in multiple paralogs revealed informative mutant phenotypes, e.g., in rice and wheat (Wu et al., 2017; Li et al., 2019). So far there have been no inflorescence phenotypes reported for A-class genes in any panicoid species, which include major cereal and energy crops. Therefore, we know little about their specific functions in regulating important agronomic traits such as flowering time and inflorescence determinacy. In this study using *S. viridis* as a model, we characterized a loss-of-function mutant in an A-class gene, *SvFul2*, that displayed strong developmental phenotypes, which was unexpected for a single mutant allele. Our morphological and molecular analyses of the *Svful2* mutant provide insights into the roles of A-class genes in connecting flowering time and inflorescence determinacy in panicoid grasses, as well as predictions on conserved and novel regulatory interactions underlying the complex phenotypes.

### SvFul2 is necessary for proper timing of flowering and determinacy programs

Phylogenetic studies have reconstructed the evolutionary history of *AP1/FUL-like* genes in angiosperms (Litt and Irish, 2003; Preston and Kellogg, 2006; Soltis et al., 2007; Wu et al., 2017). The monocot *AP1/FUL-like* clade members evolved independently after the split of monocots and eudicots (Litt and Irish, 2003; Preston and Kellogg, 2007). In the Poaceae clade, four copies of *AP1/FUL-like* genes are derived from three duplication events in the AP1/FUL lineage. The first likely occurred during early monocot evolution, giving rise to the *FUL3* clade. The second occurred near the base of the Poaceae, which generated the *FUL1* and *FUL2* clades (Preston and Kellogg, 2006). The last duplication produced the *FUL3* and *FUL4* clades, and *FUL4* was lost in some grass species during evolution (Wu et al., 2017). Such duplication events can lead to functional redundancy and subsequent diversification. In grasses, *AP1/FUL-like* genes are expressed much earlier than in eudicots (Preston and Kellogg, 2007), and their transcripts have been detected in IM, BMs, and SMs in addition to floral organs. In several grass species studied, *FUL1/VRN1/OsMADS14* and *FUL2/OsMADS15* have redundant and/or overlapping spatiotemporal expression patterns in these three meristem types, yet show different patterns within the spikelet (Preston and Kellogg, 2007). This suggests that in certain grasses, these two genes play redundant roles during the floral transition and in SM identity, but diversified roles in floral organ identity.

Among the grasses, *AP1/FUL-like* genes have been most studied at the functional level in rice and wheat (Wu et al., 2017; Li et al., 2019), where clear functions in flowering time have been demonstrated. The role of *SvFul2* in controlling flowering time is consistent with the significant accumulation of *SvFul2* transcripts (over 100-fold change in expression in the IM) during the transition from SAM to IM. *SvFul1* and *SvFul3* were also induced during this time (Fig. 6), but fold changes were not as large as for *SvFul2*, similar to what has been shown in rice (Kobayashi et al., 2012). Based on our results and previous studies, the accumulation of *AP1/FUL-like* transcripts in the IM upon the induction of florigen is likely required for promoting the reproductive transition, and this is a conserved mechanism in grasses.

The shift from a determinate to indeterminate fate in the IM of *Svful2* mutants, which was also observed in the wheat *vrn1-null*; *ful2-null* double mutant, is reminiscent of the *tfl1* mutant phenotype in Arabidopsis (Shannon and Meeks-Wagner, 1991). Previous studies that examined spatiotemporal expression of *FUL1/VRN1/OsMADS14* and *FUL2/OsMADS15* in phylogenetically disparate grasses, showed that they are most abundantly expressed in the tip of the IM (Preston and Kellogg, 2007). In both *Svful2* and wheat *vrn1-null*; *ful2-null* mutants, significant increases in the expression of *TFL1* homologs were detected (Fig. 6; (Li et al., 2019)). These results suggest that the mechanism for controlling IM determinacy in grasses involves an antagonism between *AP1/FUL-lik*e genes and *TFL1-like* genes, as in eudicots. IM determinacy appears to be very sensitive to the activity of *AP1/FUL-lik*e genes. In wheat, complete loss of both *VRN1* and *FUL2* function leads to an indeterminate IM, while a single functional copy of *VRN1* or *FUL2* in a heterozygous state was able to recover a determinate IM. It has been proposed that indeterminate growth in the IM was derived from a determinate habit in evolution, which involved the modification and/or loss of an early common *TFL1* mechanism (Bradley et al., 1997). This hypothesis could explain this apparent sensitivity.

The strong phenotypes we observed in *S. viridis* by a single knockout of an *AP1/FUL-like* gene indicates its central role in controlling multiple developmental processes. Interestingly, the co-expression of closely-related paralogs, *SvFul1* and *SvFul3*, with *SvFul2* does not seem to provide much functional compensation, but we did see both genes up-regulated upon *SvFul2* perturbation. *SvFul2* was expressed at high levels (highest among other *AP1/FUL-like* genes) at most of the developmental stages we examined. The functional redundancy of *AP1/FUL-like* genes in grasses provides an opportunity for diversification of function, and a toolkit for fine-tuning development of desired traits.

### SvFul2 as an integrator of flowering time and inflorescence determinacy

Connections between flowering time signals and meristem determinacy pathways in the inflorescence have been highlighted in various grass species. A strong flowering signal can impose meristem determinacy when perceived by the developing inflorescence (Dixon et al., 2018). For example, wheat *Photoperiod-1* (*Ppd-1*), which functions in a photoperiod-dependent floral induction pathway, suppresses paired spikelet formation through modulation of FLOWERING LOCUS T (FT) (Boden et al., 2015). The paired spikelet phenotype is associated with enhanced indeterminacy. In maize, loss-of-function in *id1*, a key player in the floral transition, leads to complete loss of meristem determinacy; instead of floral organs, plantlets are developed from every spikelet in mutant tassels (Colasanti et al., 1998). Meristem identity genes, e.g., *AP1/FUL-like* genes, have been proposed to function downstream of the flowering signal to promote meristem determinacy and reshape inflorescence architecture (Dixon et al., 2018).

In our study, the important role of *SvFul2* in coordinating flowing time and meristem determinacy is not only supported by its strong pleiotropic phenotypes, but also reflected in our predictions of regulatory relationships between SvFUL2 and its upstream modulators and downstream targets. Several MADS-box TFs, most of which are homologs to those implicated in flowering time, were predicted to directly target *SvFul2* (Fig. 7D). Our network analysis also uncovered potential feedback regulation between SvFUL2 and SvRA2, which could point to a conserved mechanism by which flowering links to AM determinacy in grasses. *SvRa2* is the ortholog of maize *ra2* and barley *Vrs4* (*Six-rowed spike4*) (Bortiri et al., 2006; Koppolu et al., 2013). Both *ra2* and *Vrs4* function in imposing determinacy on spikelet pair meristems and triple spikelet meristems in maize and barley, respectively. Although several downstream targets of *ra2* and *Vrs4* have been identified through genetic and/or transcriptomics analyses (Bortiri et al., 2006; Bai et al., 2012; Koppolu et al., 2013; Eveland et al., 2014), upstream regulators have not been described. Unlike other genes in the RAMOSA pathway, *ra2* function is highly conserved across grasses and expresses early during AM initiation, and temporally after the expression of *AP1/Ful-like* genes (Bortiri et al., 2006; Koppolu et al., 2013; Zhu et al., 2018). In addition, localization studies have shown that *ra2* and *Ful2* are expressed in overlapping domains within BMs during early inflorescence development (Bortiri et al., 2006; Preston and Kellogg, 2007; Koppolu et al., 2013). The conserved spatiotemporal expression pattern of *ra2* is consistent with it being downstream of FUL2 to potentially coordinate the flowering signal with regulation of meristem determinacy. Further functional studies are required to determine the genetic and molecular interactions between RA2 and FUL2.

Over-expression of the maize *AP1/FUL-like* gene, *zmm28*, enhanced grain yield potential through improved photosynthetic capacity and nitrogen utilization (Wu et al., 2019). In that study, they integrated RNA-seq with ChIP-seq analyses and revealed direct targets of ZMM28, which included genes involved in photosynthesis and carbohydrate metabolism. These analyses were performed in leaf tissue. Homologs of several of these targets were differentially expressed in *Svful2* mutants, including *photosystem I light harvesting complex gene 6* (Sevir.2G22720), a gene encoding a pyruvate orthophosphate dikinase (Sevir.3G253900), and gene encoding a bZIP TF (Sevir.3G396500). Although *SvFul2* encodes a different *AP1/FUL-like* gene in a different spatiotemporal context, we also observed changes in genes associated with photosynthesis and with sugar and starch metabolism in stage 3 inflorescences where the mutant was highly indeterminate compared to wild-type. There could be common regulatory interactions between *AP1/FUL2-like* genes associated with photosynthesis, carbon allocation and sugar signals that link flowering time cues from the leaf to inflorescence architecture. We know little about the mechanisms by which sugar signals interface with development, but clear links, for example with trehalose-6-phosphate, underlie flowering time (Wahl et al., 2013) and meristem determinacy (Satoh-Nagasawa et al., 2006).

The striking phenotype displayed in loss-of-function *Svful2* mutants enables us to more clearly define molecular connections between flowering time and various aspects of inflorescence meristem determinacy. One question that comes to mind is why do the pathways regulated by *SvFul2* in *S. viridis* have fewer checks and balances in terms of functional redundancy compared to other grasses? Since *S. viridis* is an undomesticated weed, one hypothesis is that selection against indeterminacy phenotypes in inflorescences of modern cereal crop species masks the ability to recover individual functions of A-class genes at the phenotypic level. Furthermore, perhaps the phenotypes presented in *Svful2* mutants provide plasticity in *S. viridis*’s adaptability to a wide range of environmental conditions. In any case, our analyses of this mutant provide a glimpse into *AP1/FUL-like* gene function in panicoid grasses and potential regulatory interactions with known players that underlie yield potential across important cereal crops.

## Methods

### Plant materials and growth conditions

The *brl1* mutant allele was isolated from an NMU mutagenized M2 population of *S. viridis (Huang et al*., *2017)*. The mutant allele was backcrossed to the reference mutagenized line (A10.1) and selfed to generate F2 segregating populations. F4 seeds were used for phenotyping, SEM, and RNA-seq experiments. *S. viridis* plants for phenotyping were grown under either SD (12 h light/12 h dark) or LD (16 h light/8 h dark) conditions (31°C/22°C [day/night], 50% relative humidity, and light intensity of 400 µmol/sq.meter/s) in a controlled hight-light growth chamber at the Danforth Center’s growth facility. *S. viridis* plants used for SEM and RNA-seq were grown under the SD conditions.

### Scanning Electron Microscopy analysis

For SEM analysis, *brl1* mutant and wild-type inflorescence primordia were harvested from young seedlings to examine the developmental defects of mutants. Samples were fixed, hand-dissected, and dehydrated as described (Hodge and Kellogg, 2014). The dehydrated samples were critical point dried using a Tousimis Samdri-780a and imaged by a Hitachi S2600 SEM at Washington University’s Central Institute of the Deaf.

### Histology

Wild-type and mutant inflorescence primordia were harvested right after the vegetative-to-reproductive transition at 11 and 15 DAS, respectively. The samples were fixed, embedded, and sectioned as described by (Yang et al., 2018). Sections (10 µm) made with a Microm HM 355S microtome (ThermoFisher, Waltham, MA, USA) were deparaffinized, stained with eosin, and imaged with a Leica M125C LED microscope.

### Bulked Segregant Analysis

M3 mutant individuals were crossed to the A10.1 reference line and resulting F1 individuals were self-pollinated to generate segregating F2 families. The F2 individuals with mutant and wild-type phenotypes were identified, and the segregation ratio was tested by a χ2 test. DNA extracted from 30 *brl1* mutant individuals was pooled to generate a DNA library. The DNA library was made using the NEBNext Ultra DNA Library Prep Kit for Illumina (NEB), size selected for inserts of 500 to 600 bp, and sequenced using 100bp single-end using standard Illumina protocols on Illumina Hi-Seq 2500 platform at the University of Illinois, Urbana-Champaign W.M. Keck sequencing facility. Read mapping and SNP calling were performed as described (Huang et al., 2017).

### Phylogenetic analysis

The coding sequences of Arabidopsis, *S. viridis*, maize, sorghum, rice, and wheat *AP1/Ful-like* family genes were obtained from Phytozome (phytozome. jgi.doe.gov) (Supplemental Data Set 2), and aligned using ClustalW to build a maximum likelihood tree with bootstrapping (1,000 iterations) in MEGA7 (Kumar et al., 2016).

### CRISPR/Cas9 gene-editing

The genome sequence of *SvFul2* (Sevir.2G006400) was obtained from the *S. viridis* v2.1 genome (https://phytozome.jgi.doe.gov/). CRISPR-P v2.0 (Liu et al., 2017) was used to design guide (g)RNAs to minimize off-targets. Two gRNAs targeting *SvFul2* were designed at the first exon and the first intron, 133bp and 395bp downstream of the ATG start codon, respectively. Using a plant genome engineering toolkit (Čermák et al., 2017), gRNAs were combined into a level 0 construct followed by insertion into a plant transformation vector. PCR amplified fragments from pMOD_B_2303 were merged using golden-gate cloning with T7 ligase and SapI/BsmBI restriction enzymes back into the pMOD_B_2303 backbone to express the two gRNAa from the CmYLCV promoter, each flanked by a tRNA. This construct, along with pMOD_A1110 (a wheat codon-optimized Cas9 driven by the *ZmUbi1* promoter) and pMOD_C_0000 modules, were combined in a subsequent golden-gate cloning reaction with T4 ligase and AarI restriction enzyme into the pTRANS_250d plant transformation backbone. The final construct was cloned into *Agrobacterium tumefaciens* line AGL1 for callus transformation of *S. viridis* ME034 at the DDPSC Tissue Culture facility. T0 plantlets were genotyped for the presence of the selectable marker, hygromycin phosphotransferase (HPT) to validate transgenic individuals. In the T1 generation, individual plants with possible mutant phenotypes were selected and the region of the target sites was amplified using PCR and sequenced. A homozygous 540 bp deletion in the 1st exon of *SvFul2* was identified. These T1 mutants were self-pollinated to obtain T2 progeny and outcrossed to ME034 and then selfed to select Cas9-free *SvFul2_KO* plants. Primer sequences used for vector construction and genotyping are listed in Table S2.

### RNA-seq library construction, sequencing, and analysis

Poly-A^+^ RNA-seq libraries were generated from pools of hand-dissected inflorescence primordia from wild-type and *brl1* mutant seedlings. Wild-type primordia were sampled at 8, 11, and 17 DAS while, accounting for the mutant’s developmental progression, *brl1* primordia were sampled at 9, 15, and 21 DAS. For each developmental stage, four biological replicates were collected, for a total of 24 data points. RNA was extracted (PicoPure RNA isolation kit; Thermo Fisher Scientific) and subjected to library preparation from 500ng of total RNA using the NEBNext Ultra Directional RNA Library Prep Kit (Illumina), size-selected for 200bp inserts, and quantified on an Agilent bioanalyzer using a DNA 1000 chip. RNA-seq libraries were processed using an Illumina HiSeq 4000 platform at Novogene with a 150bp paired-end sequencing design. On average, for each data point ∼20 million cleaned reads were generated. RNA-seq reads were quality checked and processed using the wrapper tool Trim Galore (v0.4.4_dev) with the parameters ‘--length 100 --trim-n --illumina’. Clean reads were mapped to *S. viridis* transcriptome (Sviridis_311_v2; Phytozome v12.1, (phytozome.jgi.doe.gov)) with Salmon (0.13.0) with the parameters ‘--validateMappings --numBootstraps 30’, using as index only primary transcripts (n= 38,209). Gene normalized expression levels (Transcript Per kilobase Million, TPM) (Supplemental Data Set 3) and the count matrix for downstream analyses were determined from Salmon output files and imported in R using the Bioconductor package *tximport* (Soneson et al., 2015).

Sample variance was computed based on principal component analysis (PCA) with the function *dist* and *plotPCA* on variance stabilizing transformation (vst) scaled data. Analyses of differential expression were performed using the Bioconductor package *DESeq2* (v1.22.2) with default parameters for the Wald test. The Benjamini and Hochberg method for multiple testing correction was used to classify DEGs passing the p-value adjusted cut-off of 0.05.

For Gene Ontology enrichment analysis, we generated a refined *S. viridis* GO annotation (Supplemental Data Set 5) using the GOMAP pipeline (https://gomap-singularity.readthedocs.io) (Wimalanathan and Lawrence-Dill, 2019) this to determine overrepresentation of GO terms within gene sets with the Bioconductor package *topGO*. GO testing was performed based on the hypergeometric method.

DE genes enriched in GCN modules were obtained based on the enrichment analysis using the function *enricher* from the Bioconductor package *clusterProfiler* (Yu et al., 2012) Benjamini-Hochberg multiple test corrections.

### Weighted Gene Co-expression Network Analysis

In addition to the samples described above, we included previously described wild-type *S. viridis* inflorescence primordia samples (Zhu et al., 2018): 23 additional data points from six inflorescence stages (10, 12, 14, 15, 16, and 18 DAS). This dataset (GSE118673) was re-processed using the same methods described above and used to build a reference wild-type gene co-expression network spanning *S. viridis* inflorescence organogenesis, from the transition to reproductive phase to flower development. To reduce samples bias, we first filtered out g`enes with less than 10 counts (row sum ≤ 10), then` we calculated the Euclidean distance and Perason’s correlation among samples and removed all replicates with *rho* coefficient < 0.92 or with an Euclidean score < 0.8. Based on this, two samples were removed (8 DAS rep 4 and 17 DAS rep 3). Read counts from genes (n= 26,758) and samples (n= 33) passing the above filters were normalized with variance stabilizing transformation using the function *vst* from the Bioconductor package *DESeq2*.

A signed co-expression network was built using the *blockwiseModules* function from the *WGCNA* R package (Langfelder and Horvath, 2008) with the parameters: ‘power = 16, corType = “bicor”, minModuleSize = 30, mergeCutHeight = 0.25, maxBlockSize = 30,000, MaxPoutlier = 0.05, minModuleSize = 20’. The topological overlap matrix (TOM) was calculated from the *blockwiseModules* function using the parameter ‘TOMType = “signed”’.

The module-to-developmental stage association was conducted evaluating the significance correlation of the modules eigengene and four key developmental stages defined as: i) vegetative-to-reproductive transition (8 and 10 DAS), ii) branching (11, 12, and 14 DAS), iii) meristem determinacy (15-17 DAS) and iv) flower development (18 DAS). To conduct this analysis we created a metafile where all samples were classified according to the four key stages. The R function *cor* and *corPvalueStudent* were used to test the correlation between module eigengene and the stages.

To predict targets of *S. viridis* TFs we built a complementary network using a random forest approach with the Bioconductor package GENIE3 (Huynh-Thu et al., 2010). *S. viridis* TFs were downloaded from PlantTFDB (http://planttfdb.gao-lab.org) ((Jin et al., 2017) and overlapped with the expression matrix used in the WGCNA analysis to identify the expressed TFs in our dataset (*n* = 1,265). These TFs were used as probes to predict regulatory links between the putative targets and their expression trajectories in our dataset. We ran GENIE3 with the parameters ‘treeMethod = “RF”, nTrees = 1,000’ and putative target genes were selected with a weight cutoff ≥ 0.005. Networks were explored and plotted using the R package *iGraph*.

## Data Accessibility

Data are deposited at NCBI SRA BioProject ID PRJNA649815 and will be released upon publication.

## Supporting information

Supplemental Data Set 1

Supplemental Data Set 2

Supplemental Data Set 3

Supplemental Data Set 4

Supplemental Data Set 6

## Author Contributions

JY and ALE designed and advised the research. JY, MB, JP, AC, and HJ performed genetics and morphological experiments. EB performed GRN analyses. MB performed gene editing. JY and EB performed data analyses. JY, EB and ALE wrote the paper.

## Acknowledgements

The authors would like to thank Kevin Reilly and his team at the Integrated Plant Growth Facility (Danforth Center) for chamber maintenance and plant care and Veena Veena and her team at the Tissue Culture Facility (Danforth Center) for *S. viridis* transformation. This work was funded by the National Science Foundation Plant Genome Research Program award #IOS_1733606. JP was funded by National Science Foundation REU Site grant: Research Experiences in Plant Science at the Danforth Center (NSF-DBI-REU-1156581).

## Supplemental Data

**Supplemental Figure 1**. Additional characteristics of the *brl1* mutants (Supports Figure).

**Supplemental Figure 2**. BSA analysis of the *brl1* locus.

**Supplemental Figure 3**. Genotyping of *brl1* F_2_ seedlings using a designed dCAPS marker for the SNP located at the start codon of *SvFul2*.

**Supplemental Figure 4**. RT-PCR results showing the reduced *SvFul2* expression level in mutant inflorescence primordia compared with A10.1.

**Supplemental Figure 5**. PCR genotyping of the *Svful2_KO* CRISPR edited line.

**Supplemental Figure 6**. Dynamic expression differences of genes related to meristem maintenance, plastochron, abaxial/adaxial cell fate, and meristem determinacy between wildtype and *Svful2* mutant.

**Supplemental Figure 7**. Dendrogram representing relatedness among genes based on expression across all samples and their respective module assignments (indicated by color classification).

**Supplemental Table 1**. Phenotypic measurements of *Svful2-KO* plants.

**Supplemental Table 2**. Table of primers used in this study.

**Supplemental Data Set 1**. High-confidence SNP calls for the *brl1* mutants.

**Supplemental Data Set 2**. Alignment of coding sequences of *AP1/Ful-like* genes by ClustalW.

**Supplemental Data Set 3**. Transcript abundances (TPM) for all annotated *S. viridis* genes (v2) in the wild type compared with *Svful2* mutant inflorescence primordia.

**Supplemental Data Set 4**. Differentially expressed genes with annotation at three developmental stages determined by DESeq2.

**Supplemental Data Set 5**. GOMAP for *S. viridis*.

**Supplemental Data Set 6**. Overrepresentation of functional classes among differentially expressed genes based on Gene Ontology term enrichment.

